# Single-Molecule Super-Resolution Imaging Reveals Formation of NS2B3 Protein Clusters on Mitochondrial Network Leading to its Fragmentation during the Onset of Dengue (Denv-2) Viral Infection

**DOI:** 10.1101/2022.12.18.520514

**Authors:** Jiby M. Varghese, Prakash Joshi, S Aravinth, Partha P. Mondal

**Author notes:** Corresponding author (Partha Pratim Mondal).

## Abstract

Understanding viral proteins at single-molecule level and visualizing their dynamics in a cellular environment is challenging. This calls for sophisticated microscopy techniques (such as SMLM) and image-based study that can reveal details with the precision of a single-molecule. Specifically, NS2B3 is recognized as a critical protein complex responsible for proteolytic activity and processing of viral polyprotein during dengue type 2 (*Denv* − 2) infection. Using single-molecule-based super-resolution imaging, we study the dynamics of NS2B3 protein, its kinetics and its interaction with cell organelles. Two distinct photoactivable recombinant plasmids (mEos-NS2B3 and PAGFP - NS2B3) and a fluorescent recombinant plasmid (eGFP-NS2B3) were constructed to exemplify the role of NS2B3 protein complex. The study was conducted on NIH3T3 cells and optimized transfection protocol was developed. Studies confirmed the formation of NS2B3 clusters on the mitochondrial network. Statistical analysis of super-resolution data (images) helped determine cluster dynamics and facilitated the estimation of critical biophysical parameters (such as, cluster density, number of molecules / cluster and its spread). Results indicate an average NS2B3 cluster area / spread of ∼ 0.050 *µm*^2^ with a density of 3500*mol*.*/ µm*^2^, and an average of ∼ 120 NS2B3 molecules per cluster. In addition, regional analysis suggests a direct positive correlation between NS2B3 cluster formation (single-molecule localization microscopy study) and fragmentation of mitochondrial network (confocal microscopy study). To further exemplify, we have carried out time-lapse imaging to visualize formation of NS2B3 clusters and its dynamics. The corresponding cluster parameters (#*clusters*, #*mol/cluster* and cluster area) suggests an increase in average #*mol/cluster* and cluster area. For the first time, single-molecule-based super-resolution imaging study helped reveal the dynamics of *NS*2*B*3 clusters in a cellular system. Understanding the underlying dynamics of *NS*2*B*3 clustering at the single-molecule level may help to decipher potential drug targets and the ways to disrupt the *NS*2*B*3 clusters. The image-based study has immediate implications in the broad field of single molecule imaging, fluorescence microscopy and disease biology.

**Statement of Significance:** The arrival of single-molecule super-resolution imaging techniques and image-based study have advanced the field of cell biology, and our understanding of sub-cellular processes with single-molecule precision. Here, we report the first ever application of super-resolution imaging to visualize NS2B3 clusters in a cellular system that directly links to the fragmentation of mitochondrial network. To facilitate the study, two new photoactivable probes (*mEos* − *NS*2*B*3 and *P AGF P*− *NS*2*B*3) with key protein complex, NS2B3 of dengue virus were developed. The study involves cell transfection studies followed by confocal and single-molecule imaging. The correlative biophysical study (comprising of confocal and single-molecule imaging) and estimated cluster parameters corroborates our findings.

## I. INTRODUCTION

Super-resolution imaging is perceived as the next-generation imaging technique for cell and disease biology. Recent imaging techniques have shed critical insights (at single-molecule level) in broad field range from cell biology to virology [1] [5] [6] [2] [3] [4]. In recent years, dengue has emerged as a pandemic disease in many parts of the world, with 5.2 million dengue cases in 2019 alone and spreading over 120 countries [7]. Dengue virus is an enveloped virus that consists of capsid protein (C) and a positive single-stranded RNA molecule. The dengue genome encodes a polyprotein which, upon cellular processing, releases ten proteins, of which three are structured proteins (C, prM/M, E) and seven are non-structured proteins (NS1, NS2A, NS2B, NS3, NS4A, NS4B, and NS5) [8]. Out of these proteins, it is the non-structural proteins that are directly involved in viral replication and assembly [8] [9] [10] [11] [12]. NS2B, a non-structural protease sub-unit, promotes conformational changes in the NS3 structure [13] [14] [15]. Specifically, NS3 protein has the distinct ability to cleave parts of polyprotein and is actively involved in viral replication due to its ATPase/helicase and RNA triphosphatase domain in the C-terminal region [16] [17] [18]. Together, NS2B/NS3 (in short, NS2B3) is a highly conserved protein and forms a weak non-covalent complex. The NS2B3 complex is responsible for proteolytic activity and processing of viral polyprotein at the junctions with other non-structured proteins (NS2B-NS3, NS2A-NS2B, NS3-NS4A, NS4B-NS5) [19] [20]. The NS2B-NS3 protease is crucial in correctly processing viral polyproteins during dengue viral infection.

Like most flaviviruses, dengue virus uses the host cellular mechanism to multiply and infect other healthy cells [8] [21]. However, little is known about what transpires at the single-molecule level, their interactions, and their distribution during infection. Such information is vital to understand the underlying mechanism and selective drug targets to disrupt the onset of these processes, hoping to inhibit the chain of viral infection.

Studies over the years suggest that the NS2B3 protein complex interacts with several cellular proteins, which is essential for effective infection of dengue virus to host cell [22] [23] [24]. Dengue infection begins with virion E protein binding to the host cell surface ligands [25]. The dengue virus, up on receptor-mediated endocytosis, releases viral RNA genome into host cytosol and starts producing single viral polyprotein [26]. In conjunction with host cellular proteins, the viral proteins facilitate RNA replication, budding, and complete virion maturation [27]. Dengue virion utilizes all the host organelles involved in protein expression to maturation. At the endoplasmic reticulum (ER) membrane, the NS3 protein interacts with fatty acid synthase resulting in enhanced fatty acid biosynthesis, which increases virion packaging [28]. The virion matures in the Golgi bodies, and the Golgi network helps it release into the extracellular region [29] [30].

Recent studies reported that NS2B3 protease targets host mitochondria, which induces matrix-localized GrpE protein homolog 1 (GrpEL1) cleavage. The GrpEL1 protein, a cochaperone of the Hsp-70 protein, cleavage led to the dysfunction of the Hsp-70 protein, hampering overall protein folding. Thus, GrpE1 cleavage leads to mitochondria and mitochondrial-associated membranes (MAMs) disruption, causing mitochondrial fragmentation [31]. NS2B3 proteases induce mitochondrial disruption responsible for inducing apoptosis in human medulloblastoma cells. It also blocks RIG-1 and MDA-5 translocation to the mitochondria from the cytosol leading to suppression of the host immune response [32]. However, the exact consequences of NS2B3 protease dysfunction at the molecular level are yet to be studied. Here we studied the interaction of NS2B3 protease with the mitochondrial network and evaluated its fragmentation mechanism at the single-molecule level. Most existing studies are performed by diffraction-limited microscopy techniques that are known to give ensemble information. Hence, the dynamics and related kinetics at a single-molecule or cluster level were never revealed.

Over the last few decades, the disease biology of dengue has advanced substantially with the advance of modern microscopes and biomolecular analysis. Mitochondrial damage due to dengue infection is revealed in confocal microscopy as reported by many researchers [33] [42] [43] [44]. Dengue protease, NS2B3 interferes with mitochondrial fusion and mitochondrial dynamics by cleaving mitochondrial outer membrane protein mitofusin 1 and mitofusin 2 (MFN1/2) [33]. Specifically, mitofusins help in mitochondrial fusion by tethering adjacent mitochondria [34], and decreased mitofusin activity leads to mitochondrial fragmentation [35]. It is found that MFN1 positively regulates host antiviral signaling, and MFN2 is responsible for mitochondrial membrane potential during dengue infection [36]. The cleavage of both MFNs by dengue protease would attenuate interferon production and promotes dengue-induced cell death [33]. But still, it is not well known about the mode of attachment of NS2B3 protease complex on mitochondrial membrane and its fate after attachment. There is no information available at the single-molecule level and thus, the underlying mechanism is unknown. It requires sophisticated super-resolution microscopy and single-molecule-based biological protocols. A better understanding of the infection process may lead to precision drug targeting and selective therapy for the disease.

This report presents the first imaging based study to understand Dengue (type-2) at single molecule level using single-molecule localization microscopy (SMLM). SMLM has advantages as compared to other super-resolution microscopy techniques (such as STED, structured illumination, and its recent variants) for quantifying individual molecules and their collective dynamics. These viral proteins are known to undergo processes such as migration, cluster formation, and localization to the specific organelle. Here, we study the dynamics of NS2B3 complex post-entry in the cellular system. We investigate the collective dynamics of single-molecules by suitably fusing the NS2B3 protein with a photoactivable protein. Studying the NS2B3 interaction mechanism at the molecular level has allowed us to decipher processes both at the single-molecule and cluster levels. The biophysical parameters (cluster size, density, and copy number) related to the viral protein complex are critical in assessing the progress of the Denv-2 infection process.

## II. RESULTS

In the present study, we investigate the role of NS2B3 complex post-entry to the host cell. To accomplish this, we need appropriate fluorescent probes to localize the site or organelle of activity. Two photoactivable probes (PAGFP-NS2B3 and mEos-NS2B3) and a fluorescent probe (eGFP-NS2B3) are designed. The generation of recombinant probes eGFP-NS2B3, PAGFP-NS2B3 and mEos-NS2B3 was confirmed by sequencing. The recombinant plasmids (eGFP-NS2B3, PAGFP-NS2B3, and mEos-NS2B3) were used to transfect NIH3T3 cells (mouse embryonic fibroblast cell line) to study the interaction of dengue NS2B3 with the cellular organelles.

Fig.1 shows NIH3T3 cells transfected with eGFP-NS2B3 / mEos-NS2B3 / PAGFP-NS2B3 (individual transfections) recombinant plasmids DNA followed by the standard protocol of washing, culture, and incubation for 48 hr. Here, *eGF P* − *NS*2*B*3 is used for carrying out normal fluorescence study using confocal microscopy whereas, mEos-NS2B3 and PAGFP-NS2B3 are used for single molecule studies using super-resolution microscopy. This is largely due to the fact that photoactivable probes suits better for super-resolution microscopy. The transfection was confirmed by exposing it to blue light and observing the fluorescence in green. It is clear that the protein complexes (eGFP / PAGFP-NS2B3 / mEos-NS2B3) are expressed in the cell. A few enlarged images of individual cells are also shown in Fig. 1. To ascertain the presence of NS2B3 protein complex and its proximity to the mitocondrial network, we have carried out additional studies. The experiment involves transfecting NIH3T3 cells with *eGF P* − *NS*2*B*3 recombinant plasmid. Upon expression of the *eGF P* − *NS*2*B*3 recombinant protein, the cells were washed and labeled with MitoTracker orange (Invitrogen, USA). The specimen is then fixed following standard protocol and imaged using confocal microscopy. The individual fluorescence images and their overlay images along with the transmission image are shown in Fig. 2. The proximity of *eGF P* − *NS*2*B*3 recombinant protein and MitoTracker Orange (labeling mitochondrial network) is quite evident (see, enlarged images R1 and R2 in Fig. 2). In fact, eGFP-NS2B3 clusters (green blobs in Fig. 1) are present on the mitochondrial network (magenta color). This makes mitochondria as a target organelle during the onset of Denv-2 viral infection (48 h post-transfection) as reported in other studies [31] [33]. As a next step, we studied the effect of these NS2B3 blobs on the mitochondrial network. To do so, we carried out, the image-based analysis to understand the morphological changes on mitochondrial network. Similar studies are carried out for analyzing mitochondrial fragment in cellular system [56] [57] [59]. Here, we have carried out transfection of cells followed by staining to understand the effect of NS2B3 protease on mitochondrial network. Specifically, we have transfected the cells by eGFP-NS2B3 plasmid and upon 48 *hr* post-transfection, the cells were stained and fixed. As a control, we have carried out the same process but with out transfection. Both the cases are shown in Fig. 3. The acquired images are subjected to edge-detection / skeletonization using standard inbuilt Matlab scripts and self-developed algorithms (see, right column of Fig. 3 A,B). Visually, the presence of small segments is evident in NS2B3 transfected cell, suggesting fragmentation of mitochondrial network. This can be further confirmed by comparing it with a control experiment that shows a relatively long and connected mitochondrial network. To quantify, we carried out statistical analysis related to number of fragments per unit area, fragment length, and entropy (measure of randomness). Analysis shows that, the number of fragments / area has increased by ∼ 3.6-fold for NS2B3 transfected cells (see, Fig. 3C). Specifically, the number of mitochondrial fragments per area is found to be significantly large (27.05 *±* 0.593 for *eGF P* − *NS*2*B*3) as compared to the control (7.52 *±* 0.207). This confirms the breakdown of the mitochondrial network by the viral complex NS2B3. Moreover, we have used the Shannon entropy measure to assess randomness (distribution of fragments) in each case. For control, mitotracker stained cells were expected to preserve the connectedness in the mitochondrial network, which is quite evident for control suggesting low entropy. On the other hand, eFGP-NS2B3 transfected cells show a significantly higher entropy as compared to the control cells.

**FIG. 1:**
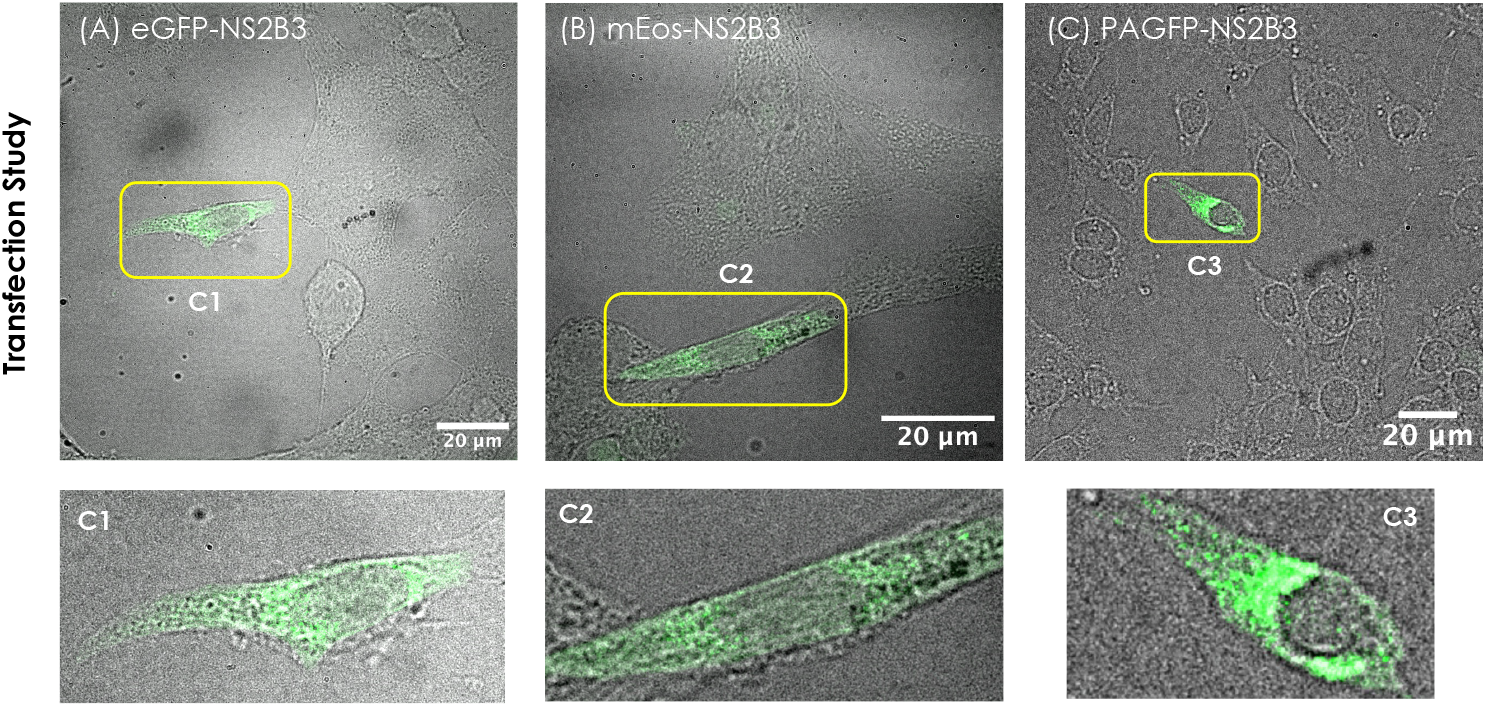
Transfection studies of NS2B3 using confocal microscope: NIH3T3 cells expressing, (A) *eGF P* − *NS*2*B*3, (B) *mEos* − *NS*2*B*3, and (C) *PAGFP* − *NS*2*B*3 protein. The transmision image is overalyed with fluorescence image. The transfected cells in each sample is marked with yellow box and their enlarged images are shown in C1, C2 and C3 respectively.

**FIG. 2:**
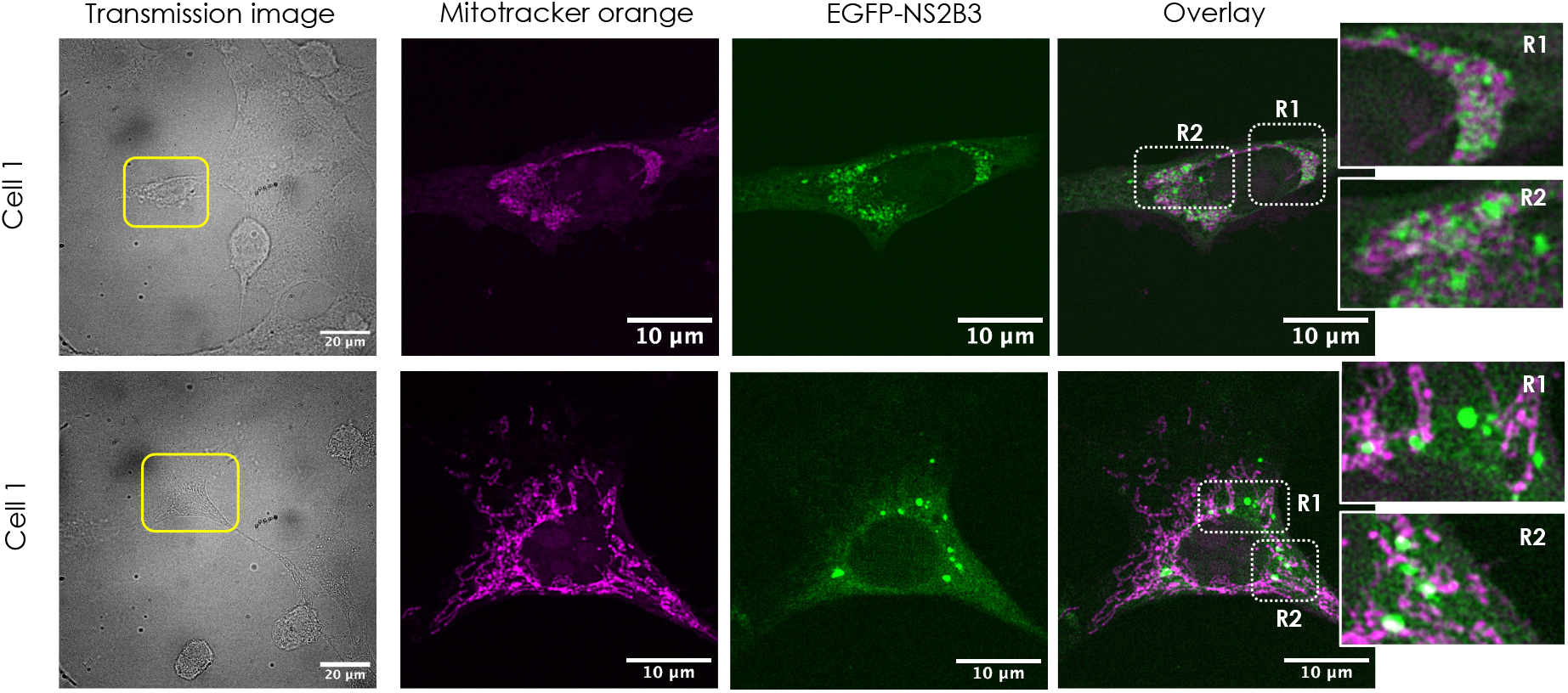
Confocal microscopy studies shows the overlay of Dengue NS2B3 protease and mitochondria : Separate and overlayed fluorescence image of the the cell sample (*EGFP* − *NS*2*B*3 transfected NIH3T3 cells (cell1 and cell2) stained with MitoTracker Orange dye). Few enlarged sections (R1 and R2) are also shown in the overlay image which clearly indicates the localization of eGFP-NS2B3 on mitochondrial network.

**FIG. 3:**
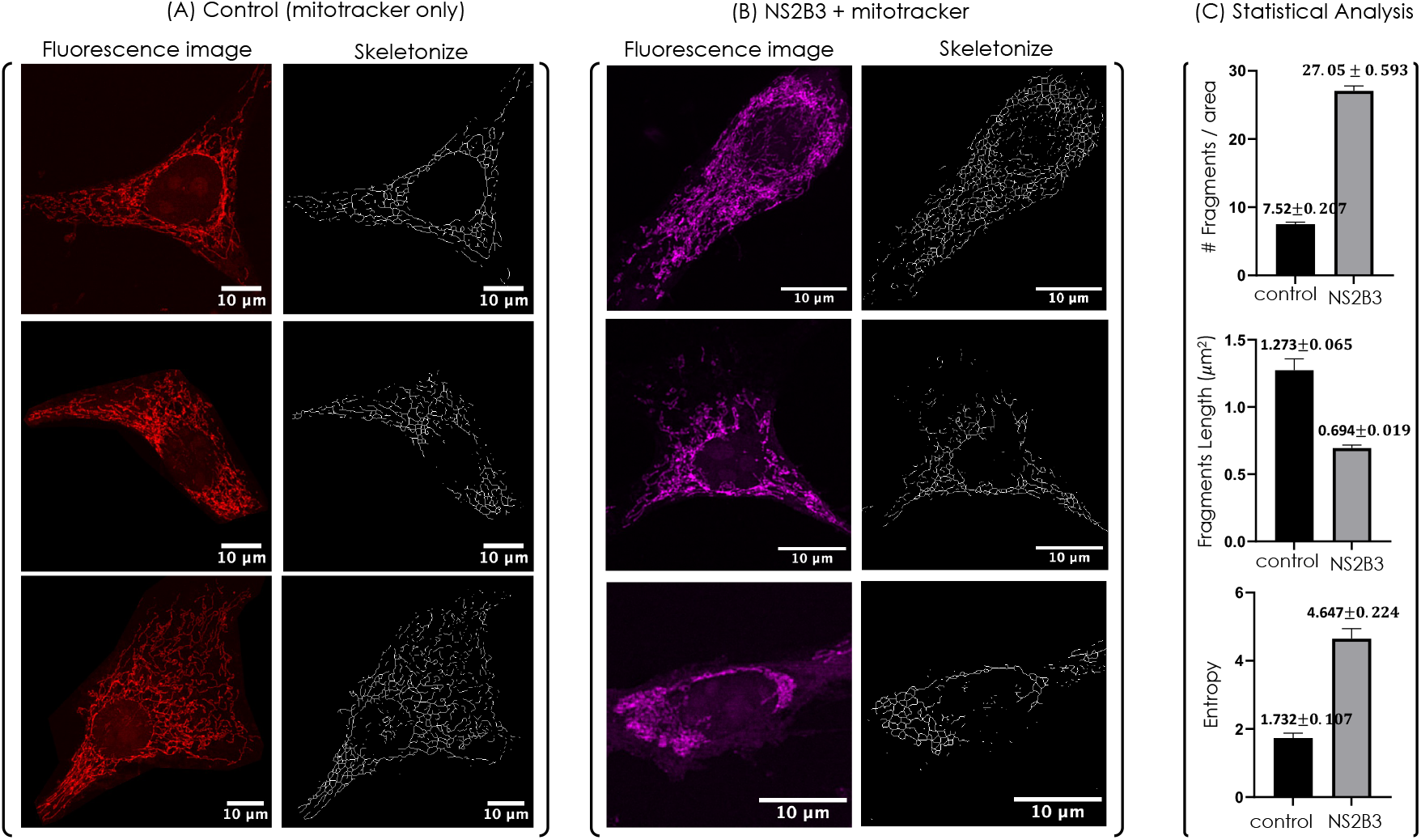
Fragmentation study showing shorter mitochondrial fragment effected by NS2B3 protease : (A) Fluorescence image for NIH3T3 cells labeled with Mitotracker Orange (Control-without NS2B3) along with skeletonized images suggesting a connected mitochondrial network. (B) Fluorescence image for NS2B3 transfected NIH3T3 cells labeled with Mitotracker Orange. Alongside skeletonized images are also shown that indicates fragmented mitochondrial network. (C) Statistical analysis for the number of fragments per unit area, fragment length and entropy are also shown. The analysis suggests a decrease in mean fragment length 0.694 *±* 0.019*µm* for NS2B3 transfected cells as compared to non-transfected cells 1.273 *±* 0.065*µm*. We have used non-parametric t-test (Mann Whitney) for statistical comparison. The corresponding statistical significance (P-values) is indicated as, * * * *p <* 0.0001. The statistical analysis presented here are based on 15 cells from 3 different experiments.

Accordingly, we set on to understand the dynamics of these viral proteins (NS2B3 complex) at the single-molecule level. We used super-resolution localization microscopy to understand the distribution of these viral proteins on mitochondria that lead to their fragmentation. The proximity of NS2B3 with mitochondria is already shown (see, Fig. 2), which is also independently reported in other studies [33] [31]. Taking this as the basis, super-resolution studies were carried out to understand the distribution of viral proteins. Here, we only focus on NS2B3 interaction and its effect on mitochondria. Fig. 4 shows super-resolved images of PAGFP-NS2B3 and mEos-NS2B3 transfected cells along with the respective transfected images. The studies were performed on fixed cells 48 h after transfection. We have used point-based clustering technique for analyzing the clusters [46]. One can readily see the appearance of single-molecule clusters indicating the formation of NS2B3 clusters. The fact that viral proteins accumulate on the mitochondrial network and form clusters suggests clustering may be necessary that ultimately leads to the fragmentation of the mitochondrial network. Corresponding localization precision analysis shows a mean value of 20 *nm*, and the resolution is calculated using FRC analysis see, methods section. It may be noted that the localization precisions for PAGFP-NS2B3 and mEos-Ns2B3 are different. This is mainly due to the photophysical properties of the respective fluorescent proteins, including the blinking period and photobleaching [47] [48] [49]. It is due to the high-resolvability of super-resolution microscopy we are able to see and analyze distinct disconnected viral clusters. Statistical analysis shows a cluster size (Area) of ≈ 0.05*µm*^2^, with a cluster density of ≈ 3500 *mol/µm*^2^ and an average of ≈ 120 *mol/cluster*.

**FIG. 4:**
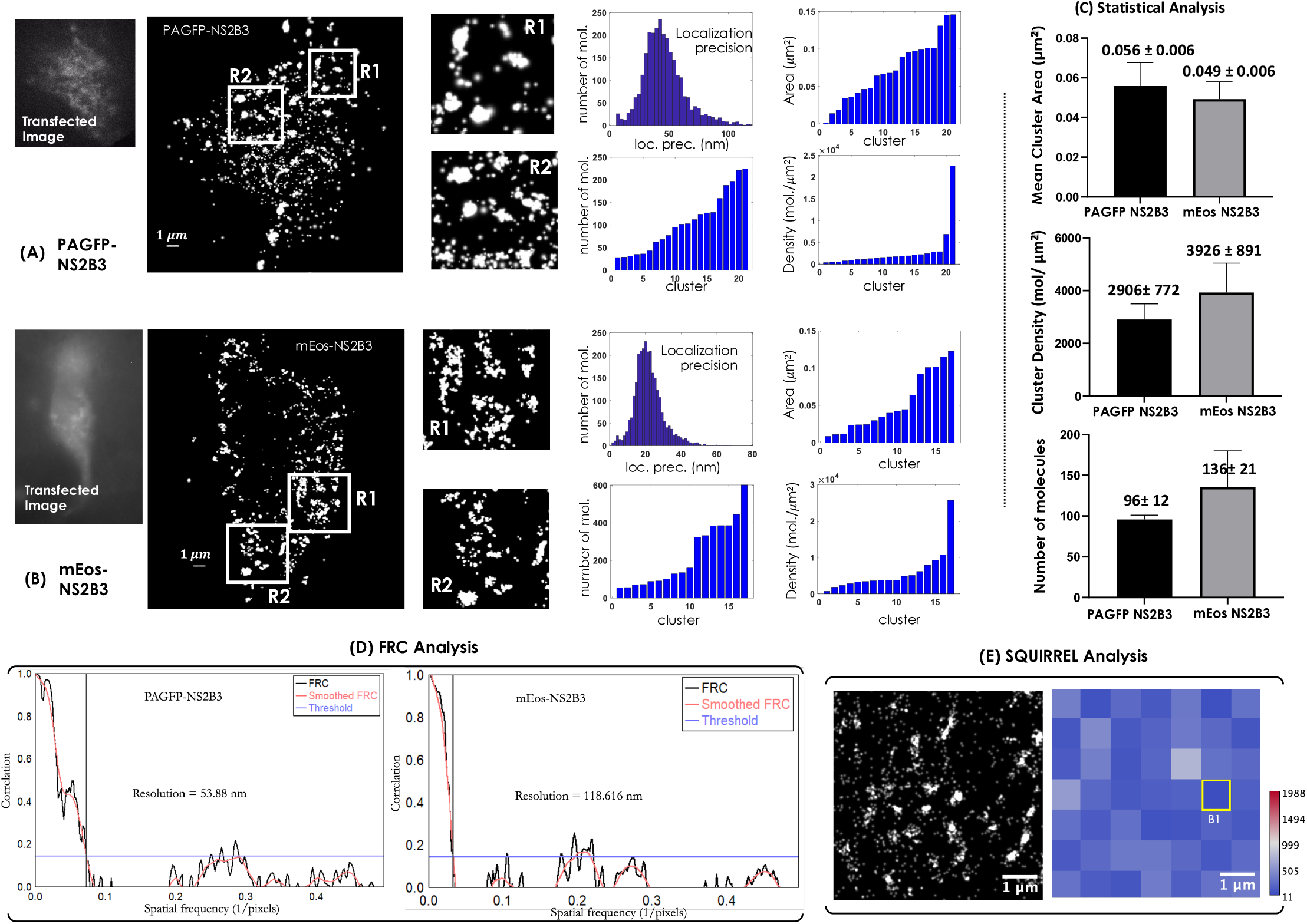
Formation of NS2B3 clusters on mitochondrial network: (A,B) Super-resolution study on the viral plasmid DNA (*PAGFP* − *NS*2*B*3 and *mEos* − *NS*2*B*3) transfected cells suggests clustering of viral proteins on mitochondrial network. This is further confirmed by enlarged section of selected regions (R1 and R2). The corresponding localization plot indicates an average localization precision of *<* 30 *nm*. (C) Statistical analysis shows that the estimated biophysical parameters have an average cluster area of 0.056 *±* 0.006 *µm*^2^ for PAGFP-NS2B3 and 0.049 *±* 0.006*µm*^2^ for mEos NS2B3 respectively. The corresponding cluster density is 2906 *±* 772 *mol/µm*^2^ and 3926 *±* 891*mol/µm*^2^ with an average number of 96 *±* 12 and 136 *±* 21 molecules per cluster for PAGFP-NS2B3 and mEos - NS2B3 respectively. The estimated cluster parameters in cells are transfected with NS2B3 gene indicate the progression of Dengue infection with single-molecule precision. (D) The Fourier Ring Correlation (FRC) analysis of NS2B3 clusters for PAGFP-NS2B3 transfected NIH3T3 cells, and mEos-NS2B3 transfected cells. (E) FRC map of reconstructed image (PAGFP-NS2B3) using SQUIRREL analysis.

Image resolution is accessed by Fourier Ring Correlation(FRC) analysis as shown in Fig. 4D. It may be noted that, the FRC curve approach zero for high spatial frequencies, and unity for low frequencies. The inverse of the spatial frequency at which the FRC curve drops below the threshold determines the image resolution. For the present analysis, a fixed threshold of 0.143 is used [61] [62]. To generate the FRC curve, we used Fiji plugin and selected 1*/*7 = 0.143 as a threshold. The corresponding Fourier Image REsolution (FIRE) number (a correlation value when the FRC curve reaches 0.143) is determined, and image resolution is calculated. Fig. 4D shows FRC for NS2B3 clusters in both PAFGP-NS2B3 and mEos-NS2B3 transfected cells, and the corresponding resolution is calculated to be 53.88 *nm* and 118.6 *nm*, respectively. The FRC map of the super-resolved image is shown in Fig. 4E. A particular portion of the FRC map (marked with yellow box, B1) is selected which shows a minimum resolution of 26*nm*.

To understand the repercussions of NS2B3 clusters on the mitochondrial network, we overlayed / superimposed super-resolved image on the confocal image (Fig. 5A). The regional analysis of biophysical parameters are carried out to determine the effect of cluster formation on the mitochondrial network locally (regions, R1-R4). So, the critical parameters (derived from confocal and super-resolution studies) such as entropy, #fragments/ area, number of clusters, and cluster density were estimated as shown in Fig. 5B and C. The visual inspection and estimated parameters show an impact on the mitochondrial dynamics related to both fission / fusion, when compared to control (non-transfected cells stained with mitotracker orange dye) (see, Fig. 5B,C). Specifically, the number of fragments/area and local entropy (regions, R1-R4) are estimated to be 0.176*/µm*^2^ and 3.89, respectively for control, wheras, in NS2B3 transfected cells, it ranges from 0.523 − 0.585 *µm*^2^ and 4.35 − 5.18, respectively. This overall suggests that NS2B3 clustering promotes mitochondrial fragmentation leading to a relatively larger number of fragments, which in turn results in an overall increase in local entropy (see Fig. 5D). Note that the cluster density is a measure of compactness of single-molecules packing per unit area. Hence, an increase in cluster density indicates more NS2B3 protease molecules packed in the given area, which gives a higher chance of fragmentation. We found both the constructs to have similar performance, although for long time imaging mEos-NS2B3 showed better performance. We used PAGFP-NS2B3 for Fig. 5 since its spectra doesn’t overlap with mitotracker orange. Overall, these critical biophysical parameters related to viral clustering on the mitochondrial network in a cellular system.

**FIG. 5:**
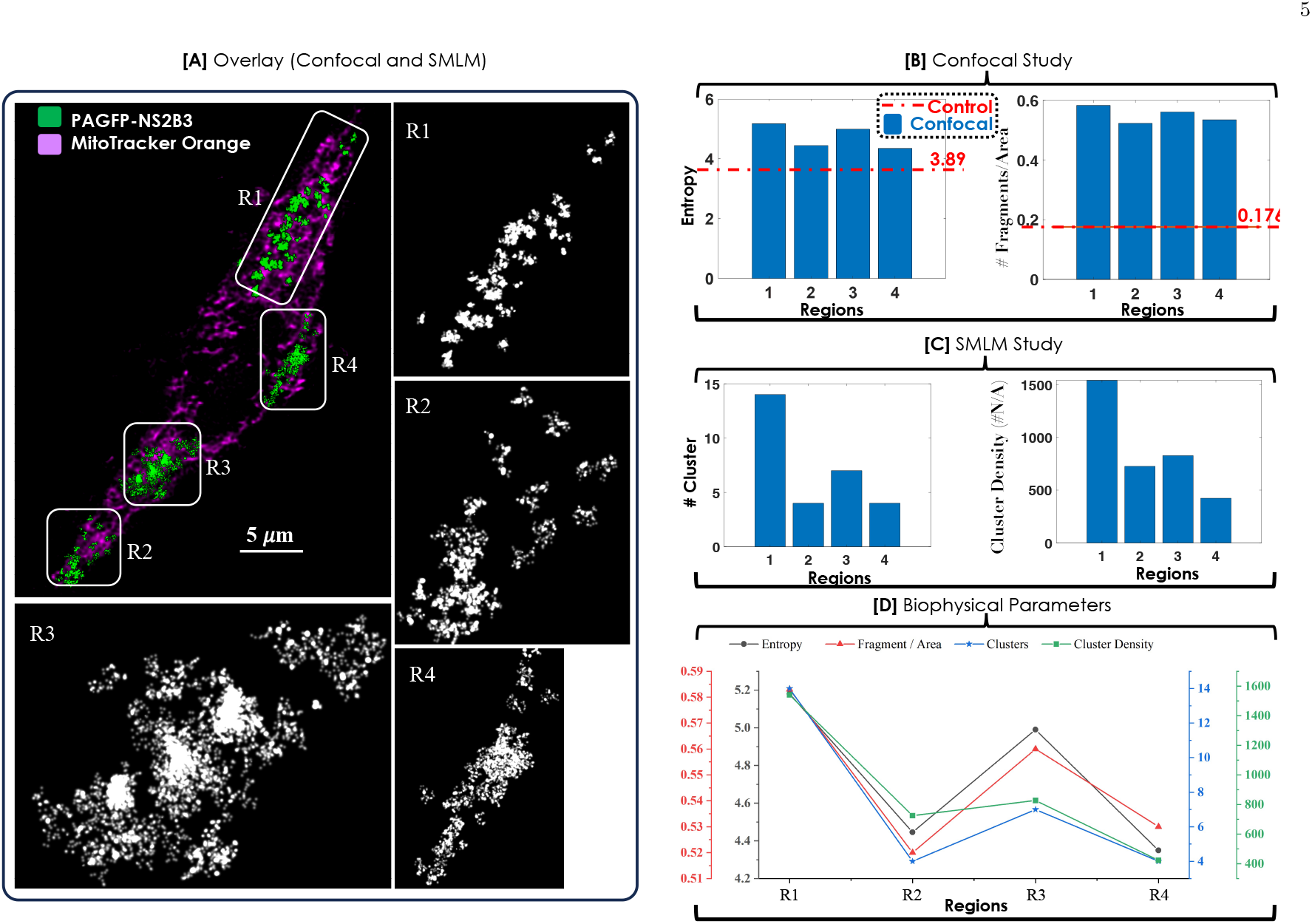
Localization of NS2B3 clusters on the site of mitochondrial fragmentation : (A) Overlaying / superimposing super-resolution data (image) on the confocal data (image) of PAGFP-NS2B3 transfected cells stained with mitotracker orange. The fragmentation of the local mitochondrial network can be noted at the site of NS2B3 clusters. This is better visualized in the enlarged view of selected regions (R1-R4). (B) Statistical analysis at the regions (R1-R4) of NS2B3 clusters indicate a proportionate increase in the number of fragments and regional entropy with respect to control (represented by red-dotted line showing an average 0.176 and 3.89, respectively). (C) Statistical analysis of single-molecule clusters suggests an increased cluster density on the site of fragmentation of the mitochondrial network. (D) Graphical representation showing the positive correlation between regional entropy, number of fragments, number of clusters, and cluster density, linking the role played by NS2B3 clusters towards the fragmentation of the mitochondrial network.

Visualizing NS2B3’s dynamics (formation/migration) provides direct evidence of clustering in a cell. To facilitate, time-lapse imaging is carried out at various time points post-transfection. This helps us to see the overall evolution of clusters, including their numbers, size, and the number of molecules per cluster. The time-lapse super-resolved image, along with the estimated biophysical parameters, are shown in Fig. 6. At each time point, the data (5000 frames) is acquired by the system and processed to reconstruct the super-resolved image (see, Fig. 6A). Subsequently, the clusters are recognized using point-based clustering algorithm and biophysical parameters are estimated (see, Fig. 6B). Specifically, we marked some of the clusters with circles (blue, green, and red). These clusters are formed dynamically over a time period of 3 *h*, showing their evolution over time. A steady increase is noted in the number of clusters (including the small ones) and the number of molecules per cluster, along with a significant increase in the cluster area (from 0.034 *µm*^2^ to 0.068 *µm*^2^). So, the time-lapse imaging and the associated analysis indicate an overall increase in the size of a few clusters over time.

**FIG. 6:**
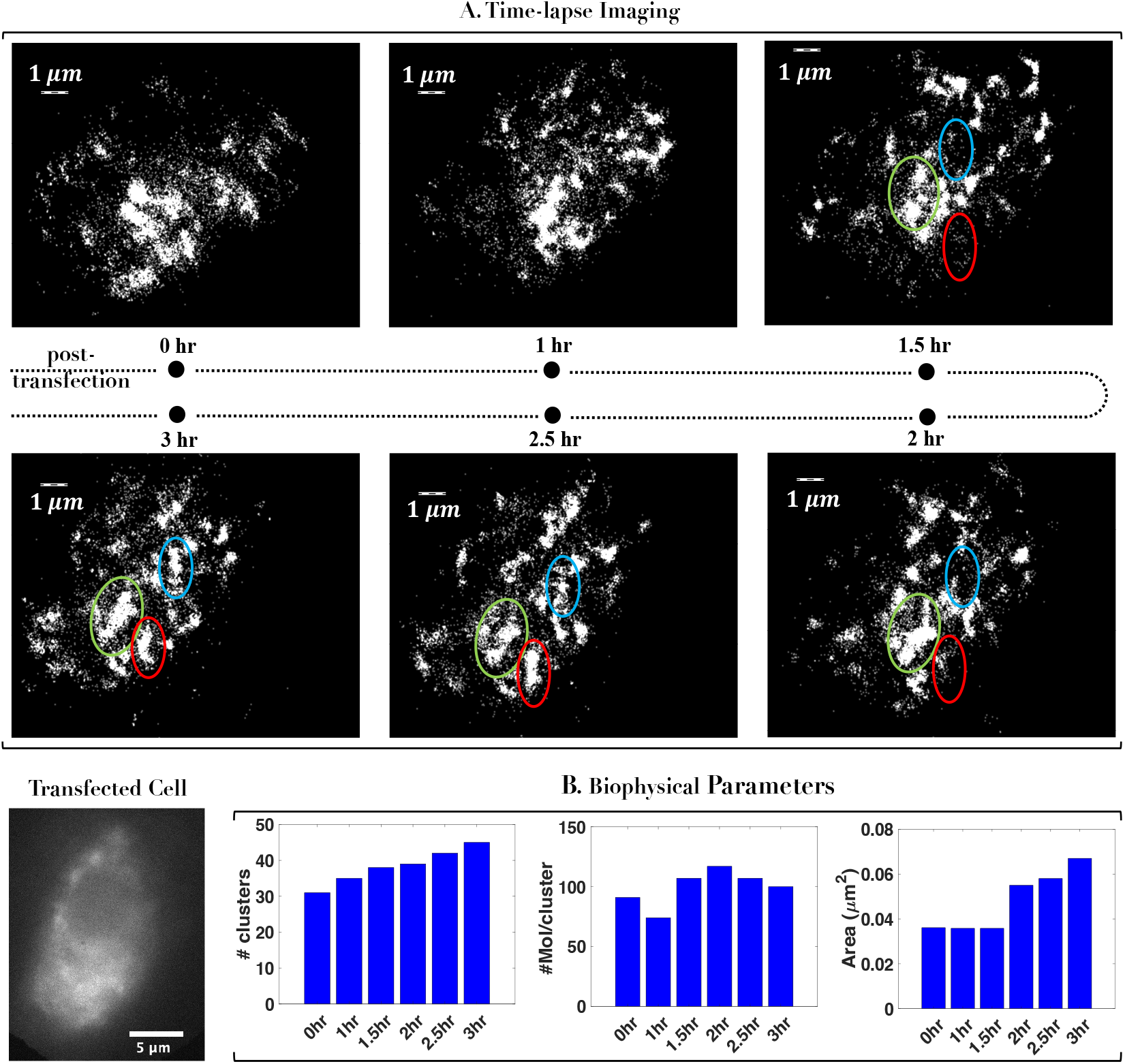
Time lapse imaging of dynamic NS2B3 clusters in live cell: (A) Raw images are collected at regular time interval (from 0 - 3 hr) and super-resolved images are reconstructed to visualize dynamically changing NS2B3 clusters. Few clusters are identified (marked by green, blue and red circles) and followed for 3 hr to visualize its formation. Alongside transfected cell image is also shown. (B) The estimated biophysical parameters (# clusters, # molecules / cluster, and cluster area) are evaluated over time indicating steady increase.

The study may provide insight to the dynamics of NS2B3 (both at ensemble and single molecule level), post entry in a cellular system that may be harnessed for targeted drug-study.

## Discussion

Understanding Dengue biophysics and pinning down the location of key protein compolexes (NS2B3) has yet to be successful. With the availability of single-molecule-based super-resolution imaging, it is now possible to decipher the process at a single-molecule level. Specifically, the key protein complex NS2B3, a protease, is known to act on the mitochondrial membrane, leading to its breakdown. Previous studies have shown NS2B3 using innate mitochondrial transport signal (MTS) to get into mitochondrial network [31]. In the present study, we report the accumulation and clustering of the NS2B3 complex on the mitochondrial membrane that leads to the fragmentation of the mitochondrial network. The in-vitro study, using purified proteins, reduced levels of matrix-localized GrpEL1, a cochaperone of mitochondrial Hsp70 (mtHsp70), noticed in NS2B3 expressing dengue infected cells is due to GrpEL1 cleavage [31]. However, in-situ details studies about the biophysical properties of NS2B3 clustering on mitochondrial membrane leading to mitochondrial fragmentation have not been studied due to the diffraction-limited resolution of existing microscopes. In this study, we used super-resolution microscopy that can decipher details at a single-molecule level to reveal the presence of NS2B3 protein clusters on the mitochondrial network, leading to its fragmentation.

Our study based on single-molecule imaging indicates that the clustering of NS2B3 single-molecules occurs at the mitochondrial network. Further, transient-transfection and protein expression studies demonstrate that a relatively large number of fragments and shorter fragment length as compared to control (see, Fig. 2 and Fig. 3). Moreover, entropy study suggests relatively more randomness in NS2B3 transfected cells. Super-resolution microscopic study and associated statistical analysis estimated the biophysical parameters (cluster size (Area) of ≈ 0.05 *µm*^2^, cluster density of ≈ 3500*mol/µm*^2^ and an average of ≈ 120 *mol*.*/cluster*) that are essential for the mitochondrial fragmentation initiation in NS2B3 transfected cells. The localization of NS2B3 clusters on the site of mitochondrial fragmentation marked the direct evidence of NS2B3 protease involvement in mitochondrial fragmentation (Figure 4). The region wise analysis of biophysical parameters like entropy, number of fragments, number of clusters and cluster density per unit area revealed the shift of mitochondrial dynamics towards fission. These parameters are critical for determining the overall progress of viral infection and, specifically, the role of the NS2B3 complex during actual Denv-2 viral infection. This is possible due to capability of super-resolution microscopy to reveal details down to single-molecule precision.

One of the findings of this study is the positive correlation between cluster formation (see, #clusters / region and cluster density plots in Fig. 5C), and fragmentation of mitochondrial network (see, entropy and #fragments plots in Fig. 5B). Previous studies have indicated that NS3 complex-induced cleavage of GrpEL1, a matrix protein is a major cause of mitochondrial matrix fragmentation [31] [50]. However, NS2B3 protease causes depletion of GrpEL1 that lead to dysfunctional mitochondria [51] [52] [52], [53] [54]. Moreover, Yu et al. (2015) have shown the association of NS2B3protease with mitochondrial membrane [33]. Time-lapse imaging is carried out to visualize the dynamics of NS2B3 molecules and the associated clustering process (see Fig. 6). The associated cluster parameters indicated an average increase in the size of a few clusters and the number of molecules per cluster over a period of 3 hr, post-transfection. Although more direct evidence may be necessary to establish the entire mechanism, the basic understanding of NS2B3 clustering can be of the therapeutic potential that may be exploited to design new drug targets or may help explore existing FDA-approved drugs.

Overall, the proposed image-based study has shown the potential to directly visualize single viral protein molecules (NS2B3), their clustering, and evolution with time. It is hoped that the proposed study may inspire the use of single-molecule techniques to investigate other viral proteins and open the door to new a kind of investigation.

## III. METHODS

### A. Recombinant Plasmids

Recombinant plasmid PAGFP-NS2B3 was generated by subcloning of NS2B3 protease from pcDNA-DENV2-NS2B3, a Gift from Alan Rothman [37] at Pst1-BamH1 site of PAGFP C1 plasmid, a gift from Jennifer Lippincott-Schwartz [38]. NS2B3 protease with PstI-BamHI overhangs were generated by PCR using NS2B3-specific oligonucleotide primers, NSPstFP with Pst 1 site as forward primer and NS-BamRP with Bam H1 site as reverse primer. Recombinant plasmid mEos NS2B3 was generated by subcloning of mEos fragment from mEos3.2-TOMM20-N-10, a gift from Michael Davidson [39] at Nhe I-Pst I site of PAGFP-NS2B3 by removing PAGFP fragment. mEos with NheI-PstI overhangs were generated by PCR using oligonucleotide primers, mEosNheFP with Nhe 1 site as forward primer mEosPstRP with Pst 1 site as reverse primer. Similary the recombinant plasmid eGFP-NS2B3 was generated by subcloning of eGFP from eGFP-N1 with Nhe I-Sac I overhangs by removing dendra 2 from Dendra 2 NS2B3 plasmid (In house plasmid where the backbone is Dendra2-Tubulin plasmid, a gift from Samuel Hess (University of Maine, USA).) The list of the sequence of primers used are given in Table 1. PCR was performed using Phusion™ High-Fidelity DNA Polymerase (Thermo Fisher Scientific, India). All restriction enzymes used were purchased from Thermo Fisher Scientific, India. Subcloning and transformation were performed using the standard protocol (Sambrook and Russel, 2001).

**TABLE 1:**
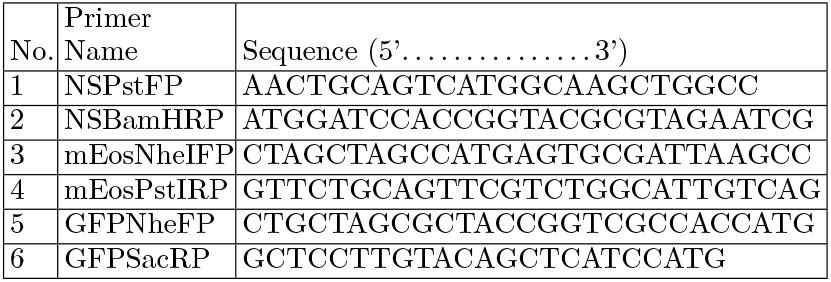
List of primers used for subcloning protein of interest (NS2B3) and the marker proteins (eGFP, PAGFP & mEos3.2).

### B. Cell Line and Transfection

The recombinant plasmids,eGFP-NS2B3, PAGFP-NS2B3, and mEos-NS2B3 were used to transfect NIH3T3 cells (mouse embryonic fibroblast cell line) to study the interaction of dengue NS2B3 with the cellular organelles. The cells used in this study were obtained from our collaborator Dr. Upendra Nongthomba (Biological Sciences, Indian Institute of Science, Bangalore, India). Transfection was performed using 1 *µg* of plasmid DNA with Lipofectamine 3000 (Invitrogen, USA) according to manufactures instructions. The cells were cultured on a coverslip (No.0) (Blue star, India) in a 35mm dish with a density of 10^5^ cells/ml. After 12h incubation cells were transfected with respective plasmids. After 48 h of post-transfection cells were washed twice with 1X PBS, and cells were incubated fixed with 3.7% (W/V) paraformaldehyde for 15 min. It was followed by washing with 1X PBS two times. Cover slip was removed from the 35mm dish and placed on a clean slide with mounting media fluorosave (Thermofisher, USA). The fixed cells were made airproof by applying nail paint on the edges for long time preservation and further investigation. The NIH 3T3 cells without transfection are considered as control. Inorder to visualize mitochondria network, the cells are stained with 175nM Mitotracker orange dye for 1 hr 15min at at 37°*C* with 5% CO2 and fixed in the same way mentioned above.

### C. Image acquisition using confocal microscopy and data processing

Fixed samples were imaged in Andor Dragonfly microscope (Bioimaging Facility, Indian Institute of Science, Bangalore, India) using 100X oil immersion objective. The cells were illuminated with a 488nm Laser, and the emission of eGFP-NS2B3, PAGFP-NS2B3 and mEos-NS2B3 was captured in the interval 502-540nm by using a band pass filter and a 40*µ*m pin hole. The images obtained from confocal microscopy were skeletonized using image J [40]. The skeletonized image was then processed using a script (inbuilt *entropy* function) written in MATLAB. The length of each fragment was measured and then the statistical analysis was performed. The entropy measures randomness of the signal in the image and it is calculated for the sample and control (mitotracker stained cells) using equation (1)

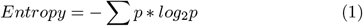

where *p* contains the normalized histogram counts from the image.

### D. Image Resolution Analysis using FRC and SQUIRREL

In optical microscopy, FRC is widely used to quantify the image resolution. Specifically, FRC is better suited for single molecule localization microscopy since it strongly depends on the localization uncertainty and density of single fluorescent labels [64] [65]. The analysis involves dividing a super-resolution image (set of single emitter localizations) into two independent sub-images (statistically independent subsets of emitters), followed by determining the statistical correlation of their Fourier transforms (in the frequency domain). Accordingly, FRC is calculated using the equation (2),

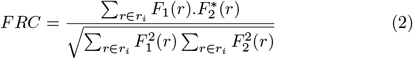

where, *F*_1_ and *F*_2_ are Fourier transform of two sub-images and *r*_*i*_ is the *i*^*th*^ frequency bin. This method gives the resolution of entire super resolution image.

Along with FRC analysis, the resolution of final super resolution image is also analyzed using an ImageJ pluggin (NanoJ-SQUIRREL) [63] and this method calculates the region wise FRC map. For the SQUIRREL analysis, the final super resolution image is transformed into two sub-images by separating the molecules equally. The sub-images are stacked and block wise FRC(Fourier Ring Correlation) is performed using NanoJ-SQUIRREL pluggin with 25 block per axis.

### E. Proximity study of mitochondria using Mitotracker orange and recombinant eGFP-NS2B3

Mitotracker orange is a dye that selectively labels mitochondria by reacting with the thiol group of cysteine residues. Then it gets accumulated in the mitochondrial matrix. Mito tracker orange (Invitrogen, USA) was dissolved in DMSO to obtain 1mM stock solution. The stock solution was diluted in DMEM to get a working concentration of 175nm, was used for staining. Cells transfected with eGFP-NS2B3 for 48 h and then stained with 175*nM* mitotracker orange followed by incubation for 1h 15 min at 37°*C* with 5% CO2. It was followed by fixing and preservation as described above. The fixed samples were imaged in confocal microscopy using two colors. We imaged eGFP-NS2B3 using laser of 488nm wavelength and the fluorescence signal is recorded with the band pass having pass band wavelength of 502-540nm. The Mito-tracker was excited using a 561nm wavelength laser and emission signal was collected with the band pass filter(572.5 - 615.5nm).

### F. Super-resolution imaging for Single-molecule analysis

Transfected cells (PAGFP-NS2B3 and mEos-NS2B3) were fixed, and super-resolution studies were carried out. We used a 405 nm laser for the activation of the samples and 488nm and 561 nm lasers were used for PAGFP-NS2B3 and mEos-NS2B3, respectively. The transfected cells were seen by 470-490 nm blue light. The system is a homemade system [45]. The fact that NS2B3 form clusters needs further investigation to determine the cluster area, number of molecules and density of clusters. The cluster analysis was done using the point-based clustering algorithm [46]. For point-clustering method, the centroid information of molecules is used. For a set of molecules to be considered as a cluster, we hace chosen a maximum Euclidean distance between two centroids as 60nm and and the minimum number of molecules as 50.

### G. Overlaying confocal and super resolution images

Cells transfected with PAGFP-NS2B3 for 48 hr and then stained with 175nM mitotracker orange followed by incubation for 1h 15 min at 37°*C* with 5%*CO*2. It was followed by fixing as mentioned above. In order to visualize mitochondrial network, the fixed samples were imaged for Mito-tracker orange using a 561nm wavelength laser in confocal microscopy. To find out the intra-cellular localization of NS2B3 clusters super resolution studies of the same cell was carried out. We used a 405 nm laser for the activation, followed by excitation at 488nm (for PAGFP fluorescent protein), for which the fluorescence signal is collected in the wavelength range 500-540nm.. The effect of NS2B3 cluster formation on mitochondrial network was analysed by comparing regional entropy, number of fragments, number of clusters and cluster density in a given region of transfected cell.

### H. Time-Lapse Imaging

To understand NS2B3 cluster dynamics, we have carried out timelapse super-resolution imaging. The transfection was performed using 1 *µg* of plasmid DNA with Lipofectamine 3000 (Invitrogen, USA). The cells were cultured on a live imaging dish (35mm dish, Mattek, USA) with a density of 30, 000 cells/ml. After 12*h* incubation cells were transfected with mEos-NS2B3 plasmid. Post 36 *h* of transfection, the cells were imaged every 30 *min* for a total duration of 3 *h*. For live cell imaging during time-lapse, the incubation temperature is maintained at 37°*C* along with 5% of CO2. The recorded images are then subjected to the reconstruction process. This is followed by cluster analysis and the determination of cluster parameters (number of clusters, number of molecules / cluster and cluster area).

### I. Statistical Analysis

All the statistical analyses were performed using GraphPad Prism 10. Data for statistical analysis were copied directly from Matlab files. Statistical significance between the test group and control were calculated by a nonparametric t-test(Mann-Whitney test). The mitochondrial branch lengths of control (stained with Mitotracker only) and sample(cells transfected with eGFP-NS2B3 and stained with Mitotracker) were considered as same (null hypothesis). From the t-test distribution table, p-value is determined for the calculated t-value. If the obtained p value is less than 0.0001, it leads to the rejection of the null hypothesis and indicates the significant difference between the control and the sample. The statistical analysis is based on a total of 15 cells from 3 separate experiments.

## Acknowledgements

The authors acknowledge ICMR postdoctoral fellowship to JMV. The authors thank Dr. Subhra Mandal (University of Nebraska - Lincoln, USA) for rigorously going through the manuscript and providing valuable inputs.

## Author Contributions

PPM and JMV conceived the idea. JMV, PJ, AS, PPM carried out the experiments. JMV, PJ, AS, PPM prepared the samples. PPM wrote the paper by taking inputs from all the authors.

## Data Availability

The data that support the findings of this study are available from the corresponding author upon request.

## Code Availability

The code that support the findings of this study are available from the corresponding author upon request.

## Disclosures

The authors declare no conflicts of interest.

## Notes

### Competing Interest Statement

The authors have declared no competing interest.

### Summary of Updates

To add more information and details.

